# Impaired light adaptation of ON-sustained ganglion cells in early diabetes is attributable to diminished dopamine D4 receptor sensitivity

**DOI:** 10.1101/2020.10.31.363564

**Authors:** Michael D. Flood, Andrea J. Wellington, Erika D. Eggers

## Abstract

Purpose: It has been known for some time that normal retinal signaling is disrupted early on in diabetes, before the onset of the vascular pathologies associated with diabetic retinopathy. There is growing evidence that levels of retinal dopamine, a neuromodulator that mediates light adaptation, may also be reduced in early diabetes. Previously, we have shown that after six weeks of diabetes in a mouse model, light adaptation is impaired at the level of ON-sustained (ON-s) ganglion cells. The purpose of this study was to determine whether changes in dopamine receptor sensitivity contribute to this dysfunction. Here we used single cell retinal patch-clamp recordings from the mouse retina to determine how activating dopamine type D4 receptors (D4Rs) changes the light-evoked and spontaneous excitatory inputs to ON-s ganglion cells, in both control and diabetic animals. We also used in-situ fluorescent hybridization to assess whether D4R expression was impacted by diabetes. We found that D4R activation had a smaller impact on light-evoked excitatory inputs to ON-s ganglion cells in diabetic retinas compared to controls. This impaired D4R signaling is not attributable to a decline in D4R expression, as we found increased D4R mRNA density in the outer plexiform layer in diabetic retinas. This suggests that the cellular machinery of dopaminergic signaling is itself disrupted in early diabetes and may be amenable to chronic dopamine supplementation therapy.

## Introduction

By 2050, it is estimated that 16 million Americans will be afflicted with diabetic retinopathy, and roughly one-fifth of these individuals will suffer vision-threatening complications of the disease (Shukla and Tripathy 2020). This presents a major challenge to healthcare in the coming decades. Although current clinical interventions largely target the vascular changes that occur with the onset of diabetic retinopathy, it has become increasingly clear that diabetes affects the neural retina long before functional changes in the retinal vasculature can be observed. Studies utilizing electroretinograms (ERGs) have identified measurable changes in retinal activity in human diabetic patients (Bearse et al. 2004; Han et al. 2004; Lynch and Abramoff 2017) who do not exhibit any clinical signs of retinopathy (i.e., vascular leakage, hard exudates, and edema). Similar deficits in ERGs can also be detected in rodent models of early diabetes (Pardue et al. 2014), and single-cell studies in these models have further identified electrical dysfunction at the single-cell level (Castilho et al. 2015a; Castilho et al. 2015b; Moore-Dotson et al. 2016; Moore-Dotson and Eggers 2019). Interestingly, multi-focal ERG studies have demonstrated that these changes in electrical activity develop asymmetrically across the retina, and that localized sites of disrupted neural activity precede and predict the sites of future vascular pathology (Fortune et al. 1999; Han et al. 2004; Harrison et al. 2011; Ng et al. 2008). Thus, it is highly likely that the neuronal changes in retinal activity which are occurring early on in diabetes are involved in the progression of DR.

In a recent study from our lab we showed that the ability of ON-sustained (ON-s) ganglion cells to adapt to increased background light was impacted by 6 weeks of diabetes (Flood et al. 2020). The retina’s ability to adapt to background light, a process known as light adaptation, allows us to see under both very bright and dim conditions and is thus crucial to normal visual function (Jackson et al. 2012; Roy and Field 2019; Witkovsky 2004). In the healthy retina, light adaptation is thought to be mediated by dopamine, a catecholamine produced and released by a single population of retinal amacrine cells, appropriately termed dopaminergic amacrine cells (Witkovsky et al. 2005). When the retina is exposed to increased light levels dopamine is released by dopaminergic amacrine cells into the retinal space (Mills et al. 2007; Perez-Fernandez et al. 2019), where it mediates its effects in a paracrine manner by binding to D1, D2, and D4 receptors located on the various retinal neuronal subtypes (Cohen et al. 1992; Derouiche and Asan 1999; Farshi et al. 2016; Klitten et al. 2008; Li et al. 2013; Veruki and Wassle 1996). There is growing evidence that suggests this dopaminergic signaling is affected in early diabetes (Kim et al. 2018; Lahouaoui et al. 2016; Tian et al. 2015; Vancura et al. 2016), and could contribute to the changes in light adaptation that we observed. However, it is currently unknown whether this disruption is attributable to changes in downstream targets of dopamine, diminished release of dopamine by dopaminergic amacrine cells, or both.

Here, we sought to determine whether dopamine sensitivity is affected in early diabetes. We examined this question in the context of ON-s ganglion cells, as we previously demonstrated impaired light adaptation in this neural population. We measured the changes induced by a dopamine D4 receptor agonist upon light-evoked and spontaneous excitatory currents in control and diabetic cells, to identify any impairment in dopaminergic signaling. In addition, we quantified the levels of dopamine receptor mRNA in control and diabetic retinas, to assess whether changes in receptor expression were responsible for dopaminergic dysfunction.

## Methods

### Animals

Animal protocols conformed with the ARVO Statement for the Use of Animals in Ophthalmic and Visual Research and were approved by the University of Arizona Institutional Animal Care and Use Committee. Experiments used C57BL/6J male mice (Jackson Laboratories, Bar Harbor, ME, USA) that were housed in the University of Arizona animal facility and given the National Institutes of Health-31 rodent diet food and water ad libitum.

Five-week-old mice were fasted for 4 hours and injected intraperitoneally with either streptozotocin (STZ, Sigma-Aldrich Corp., St. Louis, MO, USA; 75 mg/kg body weight) dissolved in 0.01 M pH 4.5 citrate buffer or citrate buffer vehicle for three consecutive days. Body weight and urine glucose were monitored weekly. Six weeks after the first injection, mice were fasted for 4 hours and blood glucose was measured (OneTouch UltraMini; LifeScan, Milpitas, CA, USA). STZ-injected animals with blood glucose ≤ 200 mg/dL and control animals with blood glucose ≥ 200 mg/dL were eliminated from the study. Fasting blood glucose was 144 ± 3 mg/dL (n=32 mice) for control mice and 418 ± 14 mg/dL (n = 29 mice; P < 0.001) for STZ-treated mice. Body weights of control and diabetic mice were 24.9 ± 0.3 and 20.6 ± 0.4 g (p < 0.001), respectively.

### Retinal slice and whole mount preparation

As previously described (Eggers et al. 2013), at 11 weeks of age mice were euthanized using carbon dioxide. The eyes were enucleated, the cornea and lens removed, and the eyecup was incubated in cold extracellular solution (see Solutions and drugs) with 800 U/ml of hyaluronidase for 20 min. The eyecup was washed with cold extracellular solution, and the retina was removed. For slice preparation the retina was trimmed into an approximate rectangle and mounted onto 0.45-μm nitrocellulose filter paper (Millipore, Billerica, MA). The filter paper containing the retina was then transferred to a hand chopper, sliced into 250-μm-thick slices, rotated 90°, and mounted onto glass coverslips using vacuum grease. Retinal whole mounts were prepared according to the methods outlined by Ivanova et al. (2013). Briefly, a hydrophilized cell culture filter insert (Millipore Sigma, Burlington MA) was trimmed down with a Dremel tool to a height of ~1mm. The retina was cut into 4 equal quadrants and mounted onto our trimmed cell culture insert as needed, one quadrant per ganglion cell experiment. Quadrants were mounted photoreceptor side down by applying slight negative pressure with a custom-trimmed 1 mL syringe. All dissections and light response recording procedures were performed under infrared illumination to preserve the light sensitivity of our preparations.

### Solutions and drug

Extracellular solution used as a control bath for dissection and whole cell recordings was bubbled with a mixture of 95% O2-5% CO2, which set the pH to ~7.4 and contained the following (in mM): 125.00 NaCl, 2.50 KCl, 1.00 MgCl2, 1.25 NaH2PO4, 20.00 glucose, 26.00 NaHCO3, and 2.00 CaCl2. The intracellular solution in the recording pipette used for monitoring inhibitory rod bipolar cell currents contained the following (in mM): 120.00 CsOH, 120.00 gluconic acid, 1.00 MgCl2, 10.00 HEPES, 10.00 EGTA, 10.00 tetraethylammonium-Cl, 10.00 phosphocreatine-Na2, 4.00 Mg-ATP, 0.50 Na-GTP, and 0.1% sulforhodamine-B dissolved in water and was adjusted to pH 7.2 with CsOH. With these concentrations, the driving force for Cl− was calculated as 60 mV in all solutions. The D4 agonist PD-168077-maleate (PD, 500 nM; Sigma) was used to selectively activate D4 receptors. This drug was diluted in extracellular solution to the given concentration and applied to the bath during the recordings by a gravity-driven superfusion system (Cell Microcontrols, Norfolk, VA) at a rate of ~1 ml/min. As DMSO was initially used to solubilize the drug, our perfusate had a final working DMSO concentration of less than 0.0025% by mass. Chemicals were purchased from Sigma-Aldrich (St. Louis, MO), unless otherwise indicated.

### Whole cell recordings

All light response recordings were first performed on retinal slices or whole-mounts in a dark-adapted state, followed by recordings performed under drug-added conditions. Retinal slices on glass coverslips or whole-mount preps were placed in a custom chamber and heated to 32° by a TC-324 temperature controller coupled to an SH-27B inline heater (Warner Instruments, Hamden CT). For D4R agonist experiments, dark-adapted light responses were measured first, followed by a 5-min incubation period with the agonist, after which light responses were again measured in the presence of continuous agonist perfusion. Whole cell voltage-clamp recordings were made from ON sustained ganglion cells in retinal slices or whole-mount preps. Light-evoked (L-) and spontaneous (s) excitatory post-synaptic currents (EPSCs) were recorded from ON ganglion cells voltage clamped at −60 mV, the reversal potential calculated for Cl^−^ currents. For all recordings, series resistance was uncompensated. Electrodes were pulled from borosilicate glass (World Precision Instruments, Sarasota, FL) using a P97 Flaming/Brown puller (Sutter Instruments, Novato, CA) and had resistances of 3–7 MΩ. Liquid junction potentials of 20 mV, calculated with Clampex software (Molecular Devices, Sunnyvale, CA), were corrected for before recording. Light responses were sampled at 10 kHz and filtered at 6 kHz with the four-pole 165-Bessel filter on a Multi-Clamp 700B patch-clamp amplifier (Molecular Devices) before being digitized with a Digidata 1140 data acquisition system (Molecular Devices) and Clampex software. Confirmation of ganglion cell morphology and presence of an axon was done at the end of each recording using an Intensilight fluorescence lamp to visualize sulforhodamine-B fluorescence and images were captured with a Digitalsight camera controlled by Elements software (Nikon Instruments, Tokyo, Japan).

### Light stimuli

Full-field light stimuli were evoked with a light-emitting diode (LED; HLMP-3950, λpeak = 525 nm; Agilent, Palo Alto, CA) that was calibrated with an S471 optometer (Gamma Scientific, San Diego, CA) and projected through the camera port of the microscope onto the stage via a 4x objective. The stimulus intensities were chosen to cover the mesopic range of light intensities (9.5, 95.0, 950.0, 9.5·10^3^, 9.5·10^4^, and 9.5·10^5^ photons·μm^−2^·s^−1^). These intensities were calculated to be equivalent to 4.75, 47.50, 475.00, 4.75·10^3^, 4.75·10^4^, and 4.75·10^5^ R*·rod^−1^·s^−1^, respectively (Field and Rieke 2002). Sequential light responses were recorded with a stimulating interval of 30 s. Stimulus intensities and duration (30 ms) were controlled with Clampex software by varying the current through the LED.

### Data analysis and statistics

Between two and four traces of L-EPSCs for each condition were averaged using Clampfit (Molecular Devices). The peak amplitude, charge transfer (Q), time to peak, and decay to 37% of the peak (D37) were determined. The bounds for integration used to calculate Q were marked by the times at which the longest-duration response for a cell began and when it returned to baseline, typically 1–2 s. These integration bounds were kept constant for all light responses recorded from the same cell. The time to peak was calculated as the temporal difference between stimulus onset and the response peak amplitude. Because the decay time could not be easily fitted with either a single or double exponential curve, we determined the D37 by computing the time it took for the EPSC to decline from its peak amplitude to 37% of its peak amplitude.

For L-EPSCs, data from each cell were normalized to the response recorded for that cell at the maximum intensity in dark-adapted conditions. If there was no discernable response for a given light intensity after filtering and averaging, the peak amplitude was recorded as 0, and it was excluded from our analysis of response kinetics. Comparisons between experimental conditions and luminance intensities were made with two-way ANOVA tests using the Student-Newman-Keuls (SNK) method for pairwise comparisons in SigmaPlot (Systat Software, San Jose, CA). If any data were shown to have a non-normal distribution or unequal variance, tests were repeated on the log10 values (or square root values for peak amplitudes) of data.

For spontaneous currents, events were included in the analysis if they occurred after a light response had returned to baseline but before the 1 s baseline preceding the subsequent light stimulus. Events were identified via the methodology outlined in (Andor-Ardó et al. 2012), using the Matlab code provided in this paper’s supplement. Frequency, amplitude, interevent interval (IEI), and decay constants for identified sEPSCs were calculated using custom-written Matlab (Mathworks, Natick MA) scripts. Decay constants were fit with a single exponent. Effects of treatments on sEPSCs were analyzed at the single cell level with Kolmogorov-Smirnov (K-S) tests. Amplitude, decay τ, and IEI cumulative distributions were normalized along the x-axis to the maximum value recorded for each cell, whether under dark-adapted or D4R-agonist-treated states. This was to allow for better visualization of the relative impact of D4R activation on sEPSC parameters across our cell populations. Effects of a D4 agonist on average sEPSC parameters in the same group of cells (control or diabetic) were analyzed with paired t-tests, and comparisons between different groups of cells (control vs. diabetic) were analyzed with unpaired t-tests, after normalizing each cell to its dark-adapted state. Individual cells were only included in the analysis if they had 10 or more spontaneous events per treatment condition. For all studies, differences were considered significant when P ≤ 0.05. All data are reported as means ± 95% confidence interval.

## Results

### D4R activation decreases the magnitude and delays the time to peak of L-EPSCs in control ON-s ganglion cells

To assess whether the action of D4R activation upon ON-s ganglion cell L-EPSCs was impacted after six weeks of diabetes we first performed light-response recordings in cells from control mice (Fig. 1A) and compared their L-EPSCs before and after application of a D4R agonist. To compensate for any differences in connectivity or responsivity between retinas, we compared normalized values for all of our analyses (see light response analysis in methods). We found that D4R activation significantly decreased peak amplitudes (2-way ANOVA, p<0.001, Fig. 1B) at all except the dimmest light intensity (SNK, P<0.05), significantly decreased charge transfer values (2-way ANOVA, p<0.001, Fig. 1C) at all light intensities above 9.5 photons·μm^−2^·s^−1^(SNK, p<0.05),and significantly delayed time to peak values (2-way ANOVA, p=0.013, Fig. 1D), although this difference seemed to arise from the lowest intensity analyzed, 95 (SNK, p=0.025). No change in D37 values was recorded (2-way ANOVA, p=0.149, Fig. 1E).

**Figure 1.**
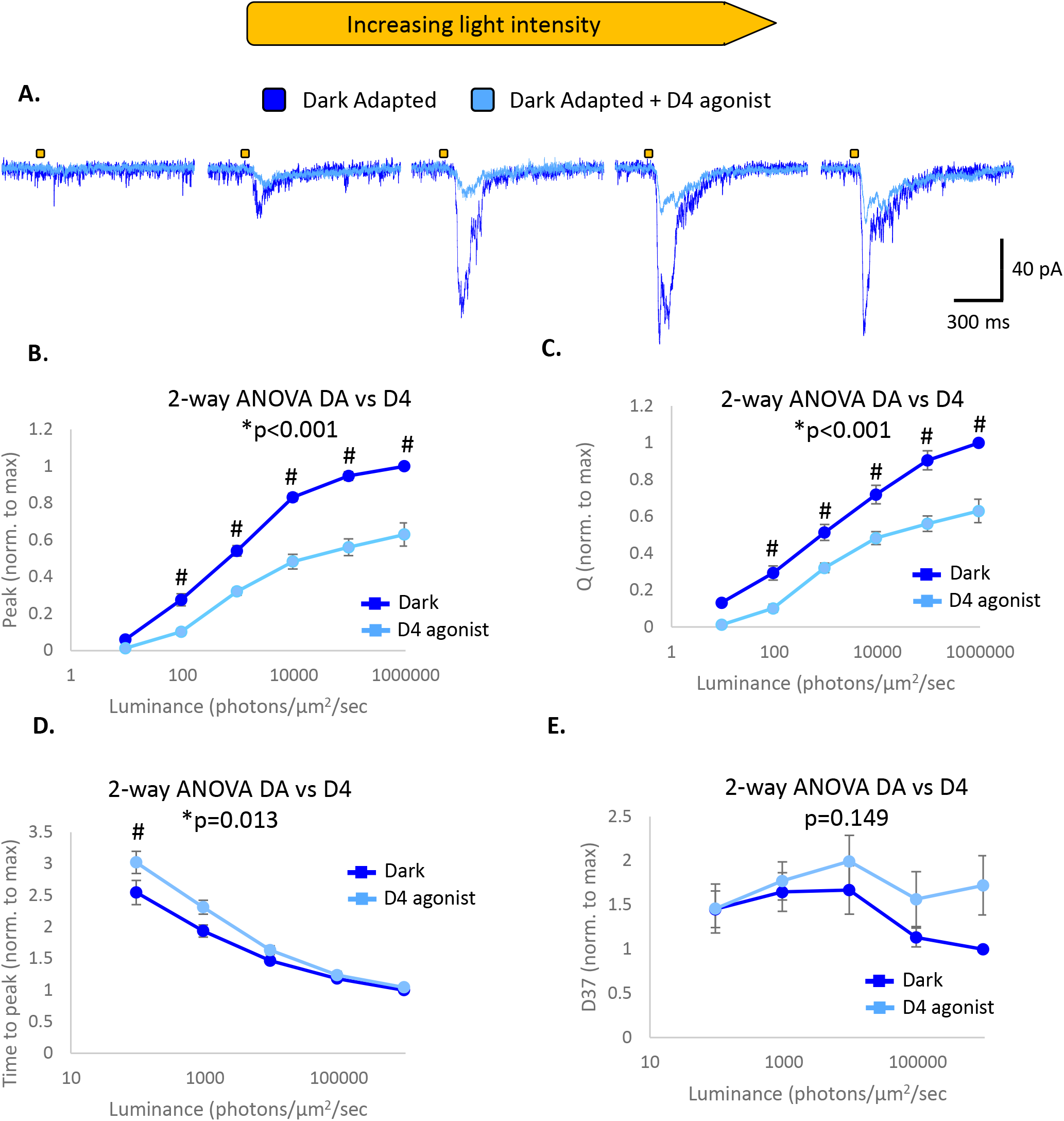
D4R activation decreases the magnitude and increases the times to peak of L-EPSCs in control ON-s GCs **A**. Example L-EPSC traces from the same cell at increasing light intensities before (dark blue) and after (light blue) application of the D4R agonist PD 168077 maleate (500 nM). The gold bars represent 30 ms light stimuli. **B-E.** Comparison of average peak amplitude (B), charge transfer (C), time to peak (D) and D37 (E) between dark-adapted and D4R-agonized conditions. All values are normalized on a cell-by-cell basis to the maximum values of each parameter recorded for each cell. Main-effects P values for two-way ANOVAs between treatment conditions are shown. * = significantly different main effect between dark adapted and agonist-treated states. # = significantly different pairwise comparison between dark-adapted and D4R activated states at specific light intensity. n = 13 cells from 10 animals.

### D4R activation decreases the magnitude of light-evoked excitatory currents in diabetic ON-s ganglion cells with no effect on kinetics

We repeated our light-response recordings in cells from diabetic mice (Fig. 2A) and compared their L-EPSCs before and after application of a D4R agonist. We found that D4R activation significantly decreased peak amplitudes (2-way ANOVA, p=0.002, Fig. 2B) at 950 photons·μm^−2^·s^−1^ (SNK, p<0.05), and significantly decreased charge transfer (2-way ANOVA, p<0.001, Fig. 2C) at 950 and 9.5*10^4^ photons·μm^−2^·s^−1^ (SNK, p<0.05). No difference was found for time to peak (2-way ANOVA, p=0..743, Fig. 2D) or D37 (2-way ANOVA, p=0.334, Fig. 2E) values. These findings suggest that even after 6 weeks of diabetes, activation of D4Rs significantly diminishes ON-s ganglion cell L-EPSCs and can continue to contribute to light adaptation.

**Figure 2.**
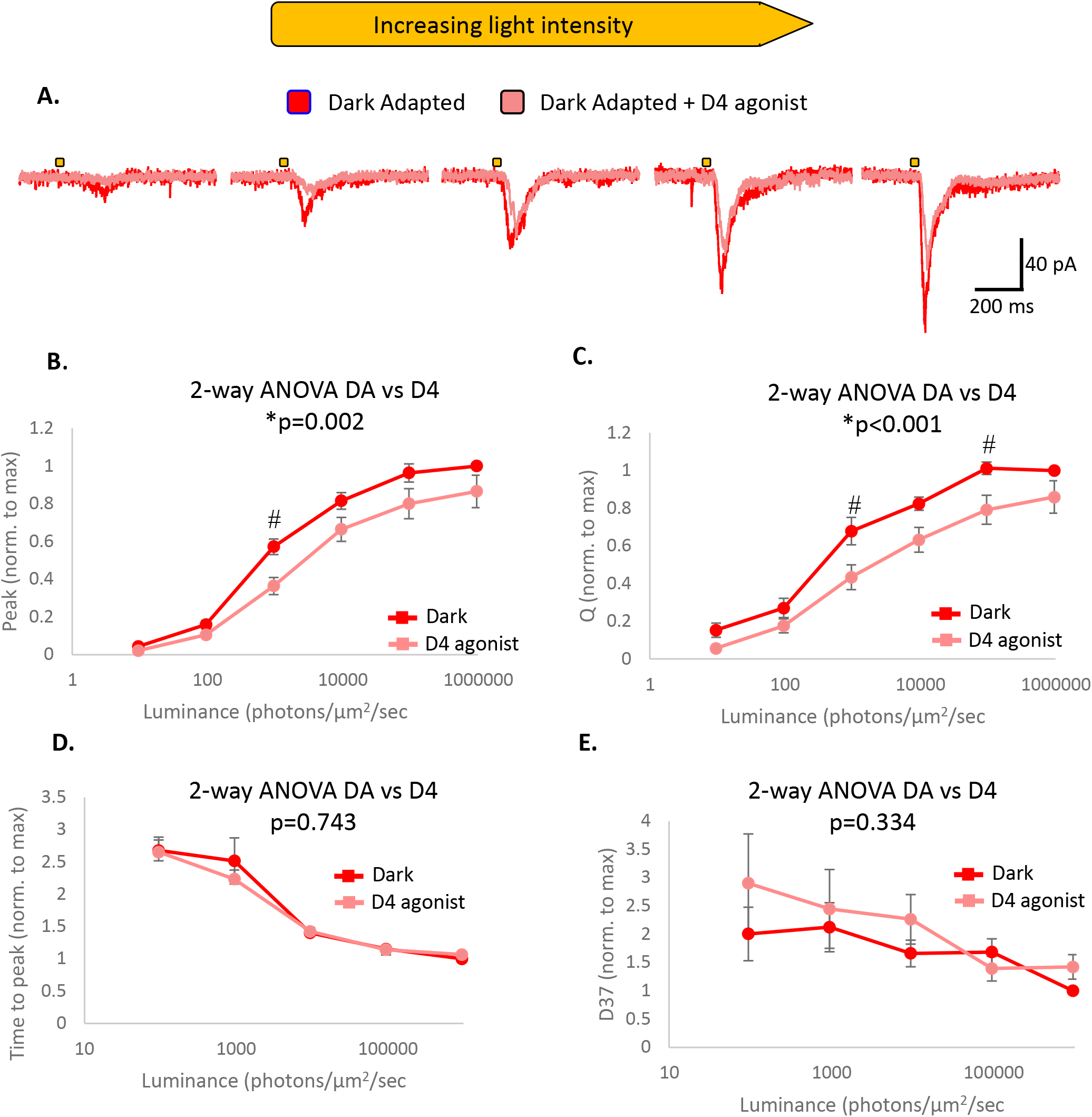
D4R activation decreases the magnitude but not kinetics of L-EPSCs in diabetic ON-s GCs **A**. Example L-EPSC traces from the same cell at increasing light intensities before (dark red) and after (light red) application of the D4R agonist PD 168077 maleate (500 nM). The gold bars represent 30 ms light stimuli. **B-E.** Comparison of average peak amplitude (B), charge transfer (C), time to peak (D) and D37 (E) between dark-adapted and D4R-agonized conditions. All values are normalized on a cell-by-cell basis to the maximum values of each parameter recorded for each cell. Main-effects P values for two-way ANOVAs between treatment conditions are shown. * = significantly different main effect between dark adapted and agonist-treated states. # = significantly different pairwise comparison between dark-adapted and D4R activated states at specific light intensity. n = 12 cells from 10 animals.

### D4R activation has a greater effect on light-evoked currents in control vs diabetic ONs ganglion cells

To assess whether D4R activation had the same magnitude of effect on L-EPSCs in control and diabetic ganglion cells, we took the values from the D4R agonist responses in figures 1 and 2 and compared them to each other (Fig. 3). These values are normalized to the response to maximum light intensity in the dark-adapted retina, so a decrease shows the effect of the D4R agonist. On average, peak amplitude values were significantly higher in diabetic than control cells, at 9.5·10^5^ photons·μm^−2^·s^−1^ and 9.5·10^6^ photons·μm^−2^·s^−1^ (2-way ANOVA, p=0.003, SNK p<0.05, Fig. 3A). Charge transfer values were significantly larger in diabetic than control cells (2-way ANOVA, p=0.002, Fig. 3B) with a significant pairwise difference at 9.5·10^5^ photons·μm^−2^·s^−1^ (SNK, p=0.011). No significant difference was found between the two in time to peak (2-way ANOVA, p=0.138, Fig. 3C) or D37 (2-way ANOVA, p=0.328, Fig. 3D). This data suggests that D4R activation does not decrease L-EPSCs to the same extent in diabetic ON-s ganglion cells as in controls.

**Figure 3.**
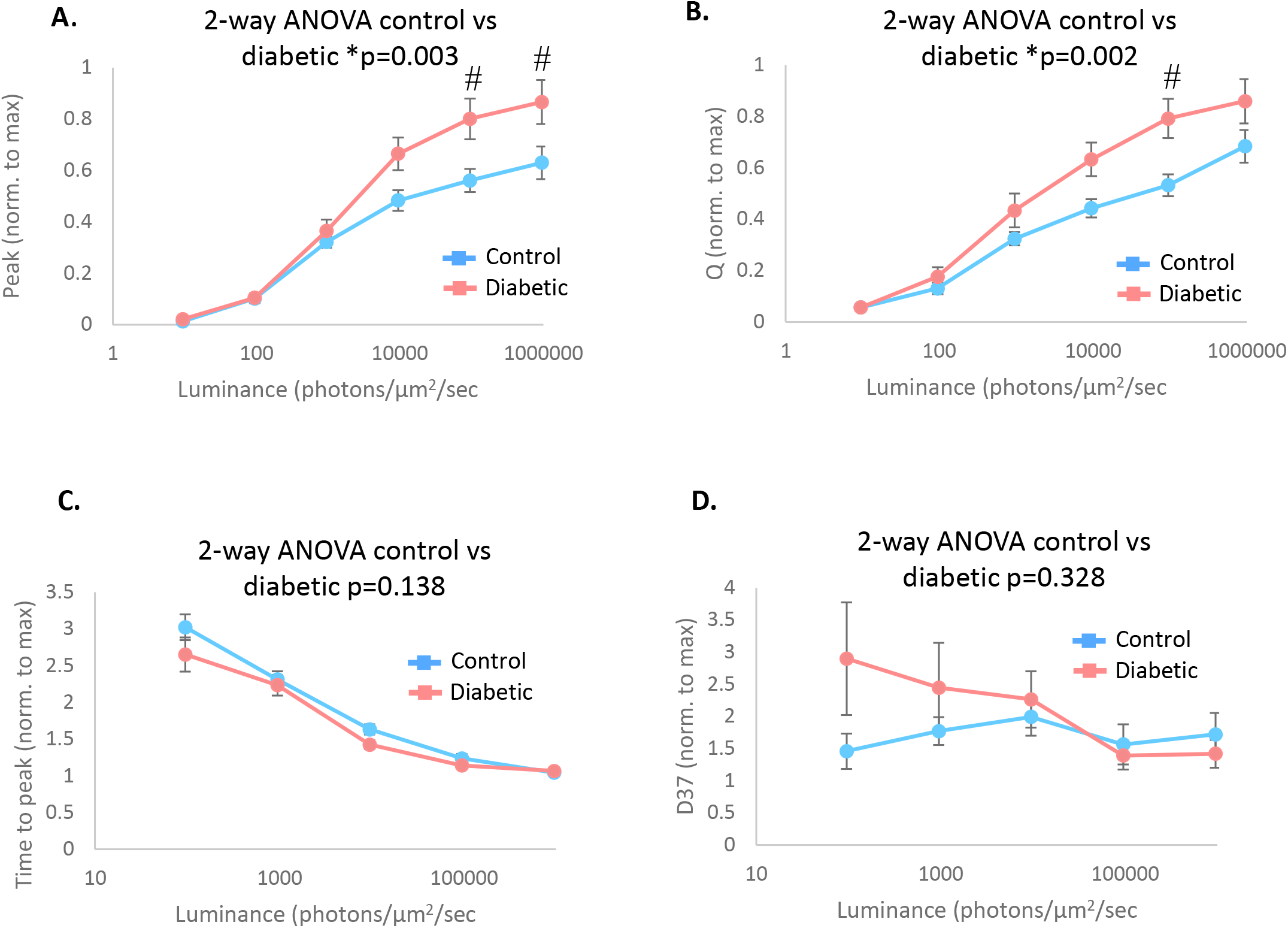
Comparison of the effects of D4R activation on L-EPSCs in control and diabetic cells. **A-D.** Comparison of normalized L-EPSC peak amplitude (A), charge transfer (B), time to peak (C), and D37 (D) values between control and diabetic cells after D4R activation. D4R activation reduced L-EPSC charge transfer and peak amplitude to a greater degree in control ON-s GCs than diabetic ones. P values for two-way ANOVAs between conditions are shown. * = significantly different main effect between control and diabetic groups. # = significantly different pairwise comparison between control and diabetic groups at a specific light intensity. Values shown here are the same as the D4 activated values from figures 1 and 2.

### Pre-synaptic effects of D4R activation on ON-s ganglion cell sEPSCs remain unperturbed after six weeks of diabetes

Previously we demonstrated that a D4R agonist decreases sEPSC frequency and amplitude in ON-s ganglion cells (Flood and Eggers 2020), suggesting changes to ON bipolar cell release and possibly post-synaptic changes in ON-s ganglion cells upon D4R activation. To help identify if the impaired effect of a D4R agonist on L-EPSCs could be attributed to changes in its post-synaptic or pre-synaptic effects, we analyzed sEPSCs from the same control and diabetic cells in which we recorded light-responses. For control ganglion cells, D4R activation decreased average sEPSC amplitude, decay tau and frequency for most cells (Fig. 4A,B), although only decay τ’s were decreased significantly (paired t-test, p=0.025). When we compared the distributions of sEPSC values in individual cells (Fig. 4C) we found that D4R activation significantly shifted sEPSC distributions towards smaller amplitudes (7/8 cells, K-S p<0.05), towards shorter decay taus (6/8 cells, K-S p<0.05) and towards longer inter-event intervals (6/8 cells, K-S p<0.05). This suggests that for most control cells D4R activation results in pre-synaptic changes to ON bipolar cell release and possibly to post-synaptic changes in ON-s ganglion cells.

**Figure 4.**
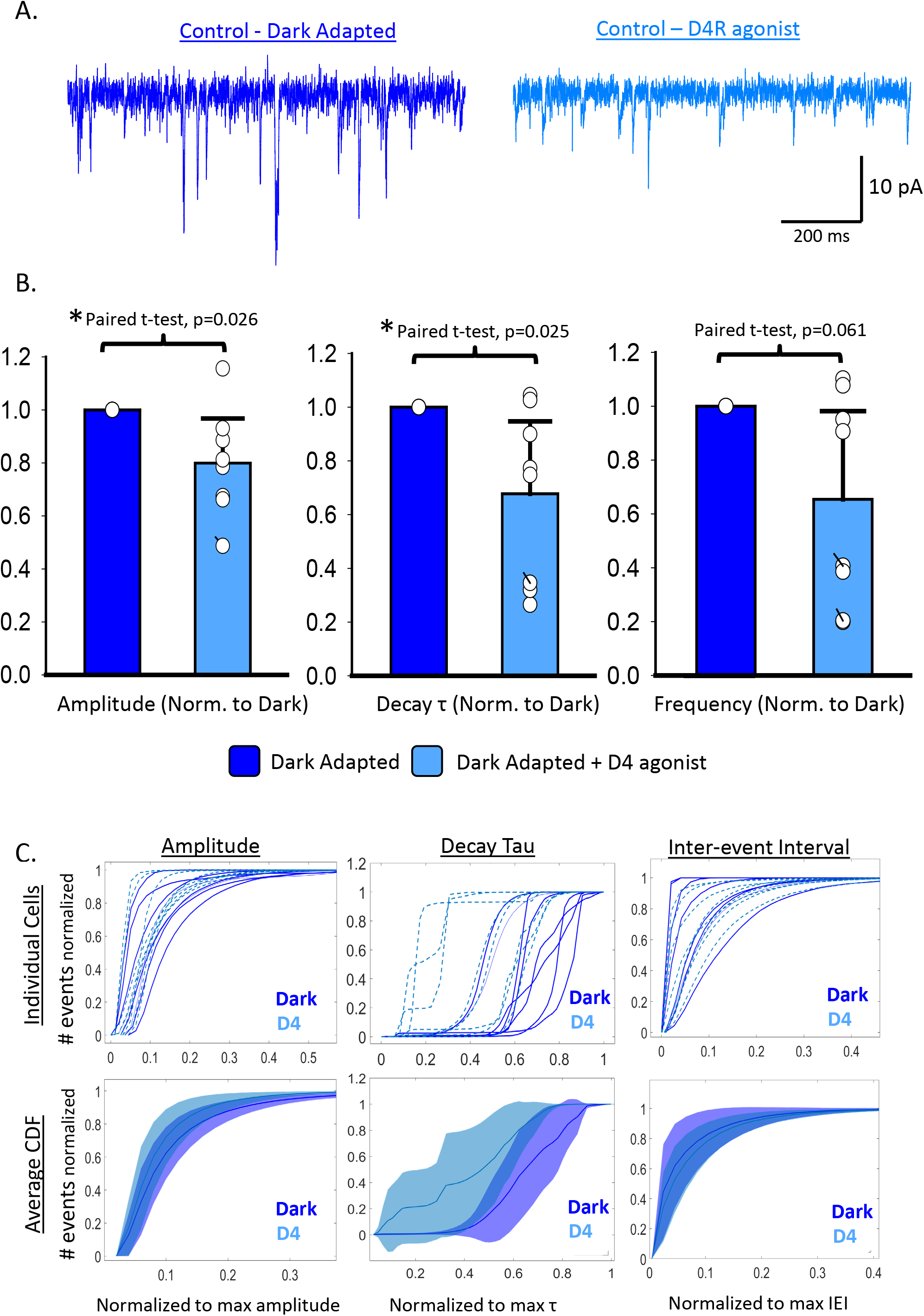
D4R activation decreases the amplitude, decay tau and frequency of sEPSCs in most, but not all control ON-s GCs. **A**. Example sEPSC traces from the same cell before (dark blue, left) and after (blue,right) application of the D4 agonist PD 168077 maleate (500 nM)**. B**. Average amplitude (left), decay τ (middle) and inter-event interval (right) values of sEPSCs before (dark blue) and after (light blue) D4R activation. Average values for individual cells are plotted as white circles. Dashed lines connect values for the same cell under dark-adapted and D4R-activated conditions. Error bars indicate average values + 95% confidence interval. **C.** Normalized cumulative histograms for sEPSC amplitudes (left) decay taus (middle) and inter-event intervals (right). **Top**. CDFs for individual cells are shown before (dark solid blue lines) and after (dashed light blue lines) D4R activation. For each CDF, the y-axis was normalized to the total number of events recorded for each condition (either dark-adapted or D4R-activated), while the x-axis was normalized to the maximum value recorded (whether under dark-adapted or D4R-activated conditions) on a cell-by-cell basis. **Bottom.** Individual CDFs were averaged for dark-adapted (dark blue) and D4R-activated (light blue) conditions. Shaded outlines represent 95% confidence intervals. n = 8 cells from 5 animals for all panels.

For diabetic ganglion cells, D4R activation significantly decreased average sEPSC amplitude (paired t-test, p=0.020), and frequency (paired t-test, p=0.032), but not decay tau (paired t-test, p=0.451, Fig. 5A-B). When we compared the distributions of sEPSC values in individual cells (Fig. 5C) we found that D4R activation significantly shifted sEPSC distributions towards smaller amplitudes (7/9 cells, K-S <0.05), towards shorter decay taus (6/9 cells, K-S <0.05) and towards longer inter-event intervals (7/9 cells, K-S <0.05). Thus, although some diabetic cells did not adhere to this pattern, the majority responded to a D4R agonist in a similar fashion as controls.

**Figure 5.**
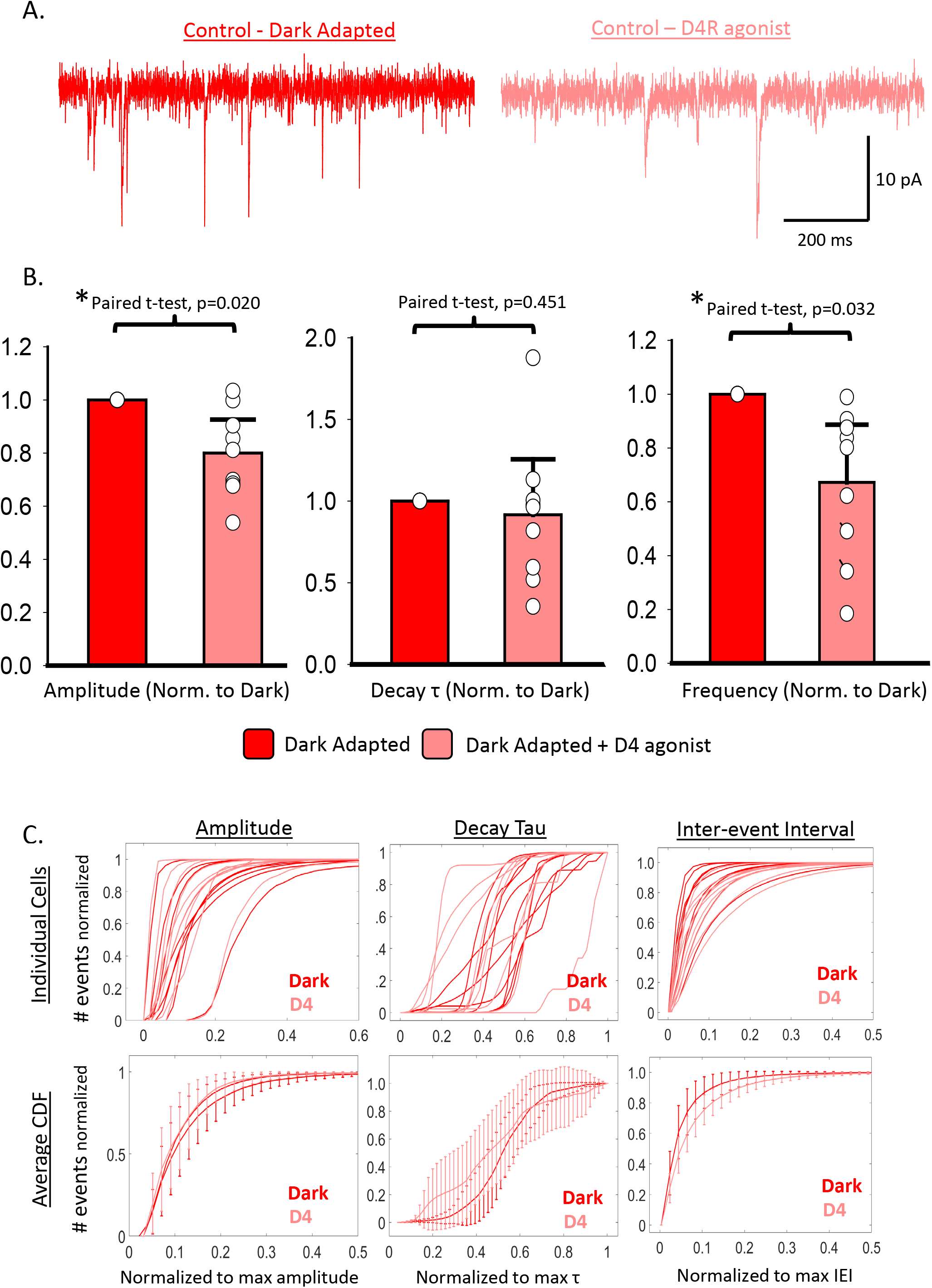
D4R activation decreases the amplitude and frequency of sEPSCs in diabetic ON-s GCs. **A**. Example sEPSC traces from the same cell before (dark blue, left) and after (blue,right) application of the D4 agonist PD 168077 maleate (500 nM)**. B**. Average amplitude (left), decay τ (middle) and inter-event interval (right) values of sEPSCs before (red) and after (pink) D4R activation. Average values for individual cells are plotted as white circles. Dashed lines connect values for the same cell under dark-adapted and D4R-activated conditions. Error bars indicate average values + 95% confidence interval. **C.** Normalized cumulative histograms for sEPSC amplitudes (left) decay taus (middle) and inter-event intervals (right). **Top**. CDFs for individual cells are shown before (solid red lines) and after (solid pink lines) D4R activation. For each CDF, the y-axis was normalized to the total number of events recorded for each condition (either dark-adapted or D4R-activated), while the x-axis was normalized to the maximum value recorded (whether under dark-adapted or D4R-activated conditions) on a cell-by-cell basis. **Bottom.** Individual CDFs were averaged for dark-adapted (red) and D4R-activated (pink) conditions. Error bars represent 95% confidence intervals. n = 9 cells from 7 animals for all panels.

To compare the magnitude of sEPSC changes induced by D4R activation directly between control and diabetic ganglion cells, we normalized average sEPSC values after D4R activation to those recorded before D4R activation on a cell-by-cell basis. When we analyzed this normalized data, we found that compared to dark-adapted values D4R activation significantly decreased average amplitude (paired t-test, p=0.0260), decay tau (paired t-test, p=0.0255), and frequency (paired t-test, p=0.0412) in control ganglion cells. For diabetic cells we still found that D4R activation significantly decreased average amplitude (paired t-test, p=0.007) and frequency (paired t-test, p=0.008) but not decay tau (paired t-test, p=0.585). This could suggest that in diabetic retinas, D4R activation fails to mediate post-synaptic changes while still mediating its normal pre-synaptic changes. When we compared the changes induced by D4R activation between control and diabetic ganglion cells directly, we found no significant difference in sEPSC amplitude (unpaired t-test, p=0.992), or frequency (unpaired t-test, p=0.913), suggesting similar degrees of modulation. Although we did expect to find a difference between control and diabetic cells for the degree of decay tau modulation, since D4R activation significantly reduced decay taus in control but not diabetic ganglion cells, we found no significant difference (unpaired t-test, p=0.229), likely due to the high variability in the data. Overall these results suggest that D4R activation still affects the output of ON cone bipolar cells to ON-s ganglion cells in diabetic retinas, but its post-synaptic actions on ON-s ganglion cells may or may not be impaired.

### D4R mRNA expression is increased after 6 weeks of diabetes

To assess whether the changes in D4R agonist sensitivity we recorded could be attributed to a decline in D4R expression, we fluorescently labeled D4R mRNA in control and diabetic retinal slices using the RNAscope system (Fig. 7A-D). Interestingly, we found that the density of D4R mRNA expression was significantly increased in the outer plexiform layer, where photoreceptor terminals and bipolar cell dendrites reside, after 6 weeks of diabetes (Fig. 7E). This finding suggests that dopamine D4Rs may actually be present at higher levels in early diabetes than in control retinas.

**Figure 6.**
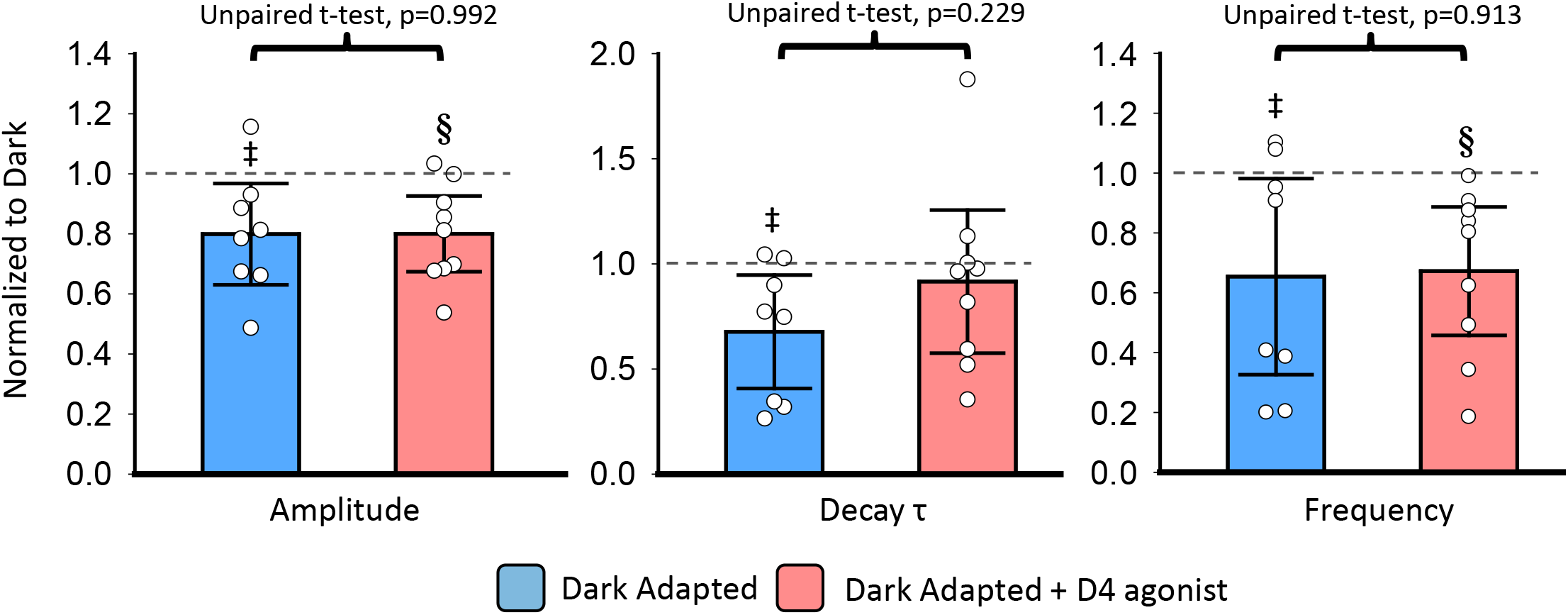
D4R activation decreases the amplitude and frequency of sEPSCs in control and diabetic ONs ganglion cells by similar degrees. Average sEPSC amplitude (left), decay τ (middle) and frequency (right) in control (light blue) and diabetic (pink) ON-s ganglion cells after D4R activation. Data was normalized to dark-adapted values on a cell-by-cell basis before averaging, and individual data points are plotted as white circles. After normalization, D4R activation significantly reduced sEPSC amplitude, decay τ and frequency in control ON-s ganglion cells. In diabetic ON-s ganglion cells, D4R activation only significantly decreased sEPSC amplitude and frequency, but not decay τ. However, when directly comparing the degree to which D4R activation reduces these parameters in control and diabetic cells, no significant difference was found. Error bars represent 95% confidence intervals. Control n = 8 cells from 5 animals and diabetic n = 9 cells from 7 animals. **‡, §**= significant difference between dark-adapted and D4R treated values in control and diabetic ganglion cells, respectively. For all tests, significance level was set to α=0.05.

**Figure 7.**
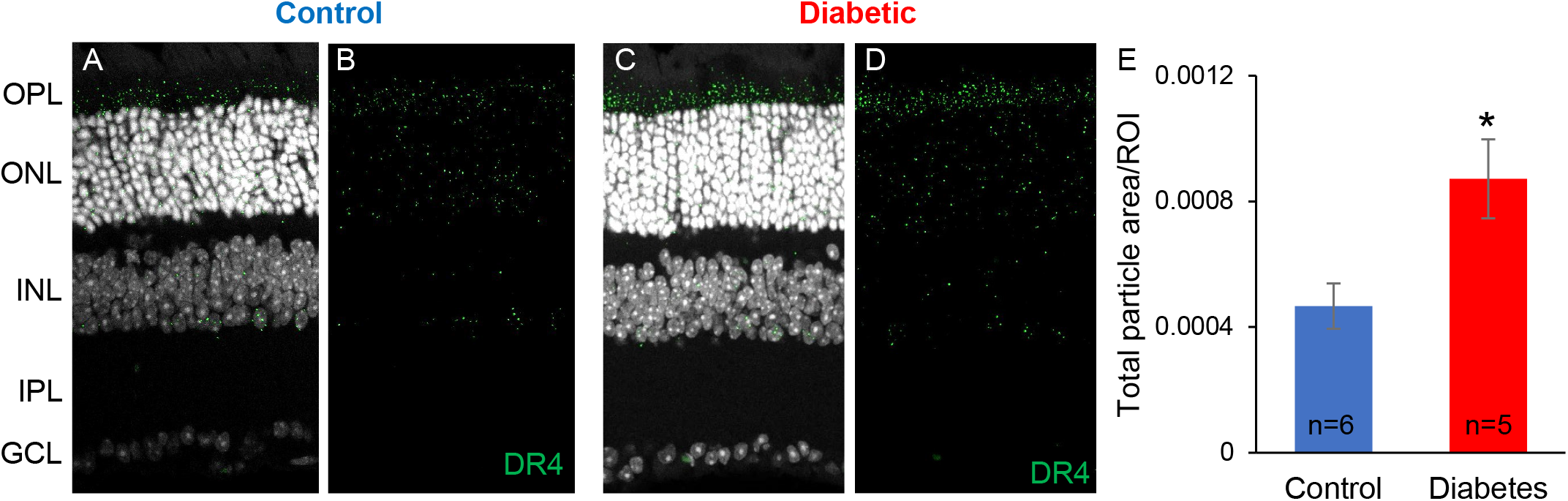
Fluorescent in-situ hybridization using the RNAscope system labeling D4R mRNA (green) and DAPI (white) in control (A,B) and diabetic (C,D) retinal slices. Total particle area was measured and normalized to the area of the outer plexiform layer (E). The diabetic retina had significantly more (t-test p<0.05) D4R mRNA area than the control. OPL = outer plexiform layer, ONL = outer nuclear layer, INL = inner nuclear layer, IPL = inner plexiform layer, GCL = ganglion cell layer.

## Discussion

Dopamine is a key neuromodulator integral to retinal light adaptation. In rodents there is good evidence to suggest that dopaminergic signaling is disrupted early in diabetes (Aung et al. 2014; Kim et al. 2018; Lahouaoui et al. 2016; Nishimura and Kuriyama 1985), though until now no specific cellular correlates of this disruption have yet been identified at this early time point. In this study we demonstrated reduced efficacy of a D4R agonist in mediating changes to retinal signaling at the ganglion cell level in diabetic mice. Additionally, we showed that this impairment in D4R signaling may occur via changes to the dopamine sensitivity of outer retinal activity as well as changes to the dopamine sensitivity of ON-s ganglion cells themselves. We found that this impaired sensitivity is likely not due to a decline in dopamine receptor expression, but may instead be caused by changes in the cellular machinery responsible for dopamine signal transduction.

### D4R activation reduces light-evoked excitation to ON-s ganglion cells in both control and diabetic retinas

Here we demonstrated that a D4R agonist reduced the size of L-EPSCs in ON-s ganglion cells in both control and diabetic animals. In the rodent retina D4 receptors are primarily expressed by photoreceptors (Cohen et al. 1992; Tran and Dickman 1992; Vuvan et al. 1993). This suggests that the main means by which D4Rs could reduce excitation at the ganglion cell level is to decrease photoreceptor light sensitivity, decrease the gain of photoreceptor output to bipolar cells, or both. In rats and mice, D4R activation has been shown to reduce cAMP levels in photoreceptors (Cohen et al. 1992; Jackson et al. 2009; Nir et al. 2002; Patel et al. 2003), thus affecting the activity of protein kinase A and its downstream actions. A recent study performed in frog retina found that rod light sensitivity was reduced by a D2/D4 agonist, possibly in a cAMP-dependent manner (Nikolaeva et al. 2019). This reduction in phototransduction would decrease output to bipolar cells for a given intensity of light stimulus. There is also evidence that D4Rs can regulate the gain of photoreceptor synapses onto bipolar cells. In mice D4Rs have been shown to negatively regulate phosphorylation of phosducin (Pozdeyev et al. 2008), a phospho-protein expressed by both rod and cone photoreceptors (Sokolov et al. 2004). In its phosphorylated state phosducin increases transmission of light responses from rod and cone photoreceptors to ON bipolar cells (Herrmann et al. 2010). This suggests that D4R activation should reduce ON bipolar cell excitation, by causing the de-phosphorylation of phosducin and thus inhibiting light response transmission. Additionally, D4R activation has been shown to decrease gap-junctional coupling between rod and cone photoreceptors in the mouse (Li et al. 2013), which further reduces the transmission of rod signals to ON cone bipolar cells (Ribelayga et al. 2008), thus decreasing overall excitation to ON cone bipolar cells. Because there is evidence that D4R expression is either unchanged (Aung et al. 2014) or increased (Fig. 7) in diabetic retinas, it follows that we witnessed decreased L-EPSCs in both control and diabetic ON-s ganglion cells after activating D4 receptors.

### D4R-mediated reduction in tonic glutamate release by ON cone bipolar cells is maintained in diabetic animals, but post-synaptic changes in ON-s ganglion cells may be impaired

ON-s ganglion cells in both control and diabetic animals exhibited reduced sEPSC amplitudes, decay taus and frequencies upon D4R activation, although decreases in decay taus were less consistent for our diabetic group. Unlike L-EPSCs, where our findings can be explained by reduced light sensitivity and signal gain, a decline in sEPSC frequency suggests a decrease in tonic glutamate release by ON bipolar cells. Since there is no evidence for D4R expression by ON bipolar cells, this reduction in sEPSCs most likely involves a D4R-mediated reduction of photoreceptor calcium concentrations (Firsov and Astakhova 2015; Ivanova et al. 2008), sustained depolarization of ON bipolar cells in response to reduced tonic glutamate release by photoreceptors, and subsequent activity-induced adaptation of bipolar cells (Snellman et al. 2008), ultimately resulting in decreased tonic glutamate release onto ON-s ganglion cells. Because we did not isolate miniature excitatory post-synaptic currents (mEPSCs) in this study, it is possible that the changes we witnessed in average sEPSC amplitude and decay tau could be explained by the decrease in event frequency, as a reduced rate of vesicle release by bipolar cells could reduce the probability of coordinated vesicle release, which in turn could affect amplitude and decay tau distributions. There is some evidence in mouse retina that sEPSCs in ganglion cells largely consists of mEPSCs (Tian et al. 1998). However, these experiments were performed at room temperature which significantly reduces the rate of spontaneous vesicle release compared to the 32°C used in our experiments. If the majority of sEPSCs we recorded do represent mEPSCs, then D4R activation seems to modify post-synaptic receptors in ON-s ganglion cells, reducing either receptor number or conductance and decreasing receptor decay times. Decay taus can also be affected by glutamate reuptake kinetics, although a previous study in salamander found that blocking these transporters had no effect on mEPSC decay taus in ganglion cells (Higgs and Lukasiewicz 1999). Several studies have reported D4R expression in some populations of ganglion cells (Cohen et al. 1992; Li et al. 2013; Van Hook et al. 2012), and there is also evidence that D2/D4 receptor agonists can modify potassium, calcium and voltage-gated sodium currents in dissociated rat ganglion cells (Ogata et al. 2012; Yin et al. 2020). Thus, it is possible that activating D4Rs on these cells could also affect glutamate-mediated currents in some manner, as D1Rs have been shown to do in OFF bipolar cells (Maguire and Werblin 1994). If this is the case, then our results would suggest some change in ganglion cell responsivity to a D4 agonist in our diabetic animals.

### D4R mediated reduction of light-evoked currents is impaired in diabetic retinas

Previous research has found that total retinal dopamine content is decreased in rodent models of early diabetes (Aung et al. 2014; Kim et al. 2018; Lahouaoui et al. 2016; Nishimura and Kuriyama 1985), and a recent study from our lab found that light adaptation was also impaired in diabetic animals (Flood et al. 2020). The simplest explanation for this previous finding would be insufficient dopamine release occurring in diabetic retinas in response to increasing light levels. However, in this study we showed that light-evoked currents were reduced by a lesser degree upon D4R agonist treatment in diabetic vs control ON-s ganglion cells (Fig. 3), suggesting a diminished capacity for diabetic cells to respond to dopamine. This means that diabetic photoreceptors and or ganglion cells either possess fewer dopamine receptors or are deficient in the cellular machinery necessary to respond to dopaminergic signaling. Our RNAscope results presented here (Fig. 7) indicate that decreased D4R expression is probably not responsible, suggesting the latter mechanism. Studies have shown that knocking out D4Rs in mouse photoreceptors reduces the absolute (Jackson et al. 2009) and circadian (Vancura et al. 2016) expression of adenylyl cyclase, the enzyme responsible for cAMP production. Application of a D4R agonist in wild-type animals increased adenylyl cyclase expression (Jackson et al. 2012). Interestingly, these results imply that the regular activation of D4Rs in photoreceptors is necessary for proper expression of the machinery that these cells require to respond to dopamine. Thus, since absolute retinal dopamine levels are decreased in early diabetes, a chronic hypoactivation of D4Rs could explain the impaired acute response to a D4 agonist that we witnessed.

Previous research has shown that retinal D4Rs are important for visual contrast sensitivity (Hwang et al. 2013; Jackson et al. 2012). One study also identified contrast sensitivity deficits in diabetic mice (Aung et al. 2014) that can be acutely resolved via injection of D4R agonists. Given these results, as well as our findings reported here, it seems likely that a deficiency in dopaminergic signaling underlies reports of impaired contrast sensitivity in diabetic human populations that lack any clinical presentation of diabetic retinopathy (Di Leo et al. 1992; Dosso et al. 1996; Ewing et al. 1998; Hyvarinen et al. 1983). In addition, there is growing evidence for D4Rs playing an important role in the circadian control of retinal metabolism (Kunst et al. 2015; Vancura et al. 2017; Vancura et al. 2016; Yujnovsky et al. 2006), as well as evidence for the disruption of these retinal circadian rhythms in early diabetes (Lahouaoui et al. 2014; 2016; Luo et al. 2019; Qi et al. 2020). If this early impairment in dopaminergic signaling is the cause for misregulated retinal metabolism, it could serve as a direct link between the visual deficits associated with early diabetes and the progression of this disease towards the more severe symptomology of diabetic retinopathy. Future work should continue to assess the link between dopamine signaling and retinal metabolism, and whether dopamine restorative therapies can provide an early intervention in diabetic models and patient populations to prevent the serious retinal complications that arise upon disease progression.

